# Fruit trichome density outweighs cuticle thickness as the dominant barrier to postharvest water loss in tomato

**DOI:** 10.64898/2026.04.09.717375

**Authors:** Xinyue Liang, Meng Li, Li Huang, Wei Zhang, Sen Zhang, Xueting Liu, Shutong Xiong, Lida Zhang, Kexuan Tang, Qian Shen

**Author notes:** **Email address:** Xinyue Liang, Meng Li, Li Huang, Wei Zhang, Sen Zhang, Xueting Liu, Shutong Xiong, Lida Zhang, Kexuan Tang.

## Abstract

Fruit cuticle thickness and biochemical composition have traditionally been regarded as the primary determinants of postharvest water loss in fleshy fruits. However, several reports indicate that some tomato mutants with thinner fruit cuticles or less cutin and waxes do not always show increased transpiration, suggesting that additional surface features influence postharvest water loss. Here, we s how that fruit trichome density is a previously underappreciated determinant of postharvest water loss in tomatoes. Using two independent mutants, *cr-slhdziv7* and *cr-slhdziv9*, which exhibit reduced fruit trichome density, we found that both mutants displayed reduced water loss rates and extended shelf life during postharvest storage despite having thinner cuticles and reduced levels of key cutin monomers. Further molecular analyses, including RNA-seq, yeast one-hybrid (Y1H), and dual-luciferase reporter assays, revealed that these two HD-ZIP IV proteins not only regulate fruit trichome formation but also directly or indirectly modulate the expression of multiple cutin biosynthesis genes. Collectively, our results demonstrate that the benefit of reducing trichome-associated microchannels can outweigh the negative effects of cuticle thinning on postharvest water loss. This study establishes fruit trichome density as an important and previously underestimated target for improving postharvest fruit quality and shelf life.

**Highlight:**

- Fruit trichome density controls tomato postharvest water loss and dominates over cuticle thickness barrier.
- Less trichomes reduce microchannels, lowering transpiration despite thin cuticles and extending tomato fruit shelf life.

## Introduction

Postharvest water loss, driven primarily by transpiration, represents a major cause of quality deterioration and economic loss in fleshy fruits globally. Since stomata are typically absent or inactive in many mature fruits, the hydrophobic cuticular layer covering the epidermis serves as the principal barrier governing water flux to the atmosphere (Fernandez-Munoz et al., 2022; Liu et al., 2023). Chemically, fruit cuticle consists of cutin, a cross-linked lipid polyester that forms the structural matrix, with diverse hydrophobic waxes embedded or deposited on its surface (Yeats and Rose, 2013). Altering the amount or architecture of cutin, or the quantity or chemistry of waxes, can significantly shift cuticular permeability. Therefore, cuticle thickness and biochemical composition of the cuticle have historically been considered the principal determinants of fruit transpiration and postharvest water loss (Lara et al., 2015; Shi et al., 2025). Defects in the cutin matrix, such as thinning or chemical alteration, often result in increased cuticular transpiration and greater susceptibility to postharvest shrinkage (Shi et al., 2013; Tafolla-Arellano et al., 2018). Conversely, thicker and well-crosslinked cuticles, on the other hand, generally confer superior barrier properties (Wu et al., 2024b). Therefore, the cuticle physicochemical properties and structural integrity thus determine the baseline for postharvest water loss and shelf-life (Fernandez-Munoz et al., 2022; Lara et al., 2019).

However, accumulating evidence has revealed that the relationship between cuticle traits and fruit water loss is more complex than previously appreciated. Several studies have reported that reducing cuticle thickness or altering its chemical composition does not invariably lead to increase in postharvest water loss in fruits. For example, in tomato, *Cutin Deficient 2* (*CD2*), which encodes an HD-ZIP IV transcription factor, plays a key role in regulating the process of cutin biosynthesis. In the *cd2* mutant plant, fruit cuticles exhibit a significant reduction in cutin content, exceeding 95% relative to the wild type (Isaacson et al., 2009). Nevertheless, the postharvest water loss and fruit phenotype analysis demonstrated that the water loss values were only marginally higher in the *cd2* mutant than in the wild type, and the phenotypes correlated with water loss appeared similar to the wild type fruits (Isaacson et al., 2009; Nadakuduti et al., 2012). It was proposed that the unaltered wax content accounted for the absence of differences in water loss; however, no consideration was given to the formation of trichomes at the fruit surface. Trichomes are epidermal hair-like structures that differ in morphology, cellular complexity, and function among species and organs, and are widely recognized as regulators of plant-environment interactions. In plants, both glandular and non-glandular trichomes can affect herbivory (Vendemiatti et al., 2024), pathogen interactions (Shen et al., 2023), and the local microclimate (An et al., 2023; Glas et al., 2012).

Previous studies have indicated that the HD-ZIP IV type genes are preferentially expressed in the outer cell layer (referred to as the L1 layer), where trichomes are initiated and developed (Chew et al., 2013). The tomato genome contains multiple *HD-ZIP IV* genes, and many of these genes have been reported to play important roles in trichome formation (Li et al., 2025; Schrick et al., 2023; Wu et al., 2024a; Wu et al., 2024c; Zocca et al., 2025). However, these studies mainly focused on trichome formation on leaves and stems (Vendemiatti et al., 2024; Xie et al., 2022; Yang et al., 2011), and with limited attention to trichomes on tomato fruit. Notably, the tomato fruits possess abundant fruit trichomes, but they are fragile and frequently broken during the harvesting and handling process. Damage and removal of tomato fruit trichomes will result in the emergence of microchannels at the base of detachment. Recent morphological and physiological studies have indicated that these fruit trichome-associated microchannels may act as potential channels for postharvest transpiration (Fich et al., 2020; Fu et al., 2025; Fu et al., 2026; Li et al., 2025; Zhang et al., 2025). Therefore, it is worthwhile to use genetic materials in which fruit trichome density was reduced to evaluate and quantitatively analyze the relative importance of trichome density versus cuticle thickness and composition for fruit water loss.

To test this, we used two tomato mutants, *cr-slhdziv7* and *cr-slhdziv9*, which exhibit substantially reduced fruit trichome density to evaluate the effects of fruit trichome density and cuticle traits on postharvest water loss. Both *cr-slhdziv7* and *cr-slhdziv9* mutants exhibited a significant reduction in postharvest water loss compared with wild type fruits. Further analysis of the mutant fruit peels revealed that their cuticles were thinner. Our data also provide molecular evidence supporting the view that *SlHDZIV7* and *SlHDZIV9* positively regulate cuticle formation. This study demonstrates that fruit trichome density is an important determinant of tomato fruit postharvest storage quality and shelf life. It can override the influence of cuticle thickness and composition on postharvest water loss. We propose that fruit trichome density represents a promising target for breeding and biotechnological strategies aimed at improving fruit shelf life.

## Materials and Methods

### Plant materials and growth conditions

The plant materials used in this study included tomato (*Solanum lycopersicum*) “Micro-Tom” and *Nicotiana benthamiana*. Both tomato and tobacco seeds were maintained in our laboratory. The *cr-slhdziv7* and *cr-slhdziv9* mutants, generated previously in a “Micro-Tom” background using CRISPR-Cas9 technology (Li 2025; Li et al., 2025), were used in this study, with “Micro-Tom” serving as the wild type (WT). Seeds were germinated on moistened filter paper at room temperature. Seedlings were then cultivated in soil on a climate-controlled artificial plant cultivation shelf (25 ± 2□, 16-h light/8-h dark photoperiod, and 50-60% humidity). The tomato fruits were tagged on the day of anthesis and harvested at 20, 30, 40 and 50 days post-anthesis (DPA) for subsequent analyses.

### Trichome quantification

Fruit trichome density changes on the epicarp of WT and mutant tomato plants were initially observed and photographed using a zoom stereomicroscope (SZ810, CNOPTEC, China). Further observation and photography were conducted using a scanning electron microscope (Hitachi, S3400 II, Japan). Pericarp tissues (20 DPA) from the same position of WT and different mutants were fixed in 2.5% glutaraldehyde solution at 4 °C for 24 h and rinsed four times in 0.1 M phosphate buffer. Subsequently, the samples were dehydrated using a gradient of increasing concentrations of ethanol (30%, 50%, 70%, 80%, 90%, 95% and 100%). The dehydrated samples were freeze-dried in a carbon dioxide critical point dryer. Dried samples were mounted on sample stubs and sputter-coated with gold in a vacuum coater (Galdon-Armero et al., 2020).

### Cuticle permeability assays

Leaf cuticle permeability was assessed by immersing detached leaves in 50 mL of 0.05% (w/v) toluidine blue (TB) solution for 2 h. Fruit cuticle permeability and trichome-associated pore quantification were of fruit trichome-associated pores were measured as described previously (Fich et al., 2020) with minor modifications. The WT, *cr*□*slhdziv7* and *cr*□*slhdziv9* fruits at the 20 DPA and 50 DPA (with rubbed and unrubbed fruit trichomes) were soaked in a 0.05% (w/v) TB solution for 12 h. After treatment, leaves and fruits were gently rinsed with distilled water to remove excess dye.

### Histological staining and cuticle thickness measurement

For fruit cuticle thickness measurements, pericarp at four developmental stages (20DPA, 30DPA, 40DPA and 50DPA) were collected from WT, *cr*□*slhdziv7* and *cr*□*slhdziv9* lines. For each line, four independent pericarp equators were cut into 6-mm cubes and fixed in Formalin-Acetic Acid-Alcohol buffer (Formaldehyde (37%): glacial acetic acid: 50% ethanol = 1:1:18). After dehydration with sucrose solution, samples were embedded in Optimal Cutting Temperature Compound (OCT). Sections (8 μm) were prepared using a freezing microtome, stained with Oil Red O and observed under a light microscopy. The cuticle thickness (Z1) and cuticle permeation thickness (Z2) were measured using ImageJ software. The statistics data of cuticle thickness and cuticle permeation thickness come from 20 slices of 4 fruits. 2-3 positions were selected for each slice to measure the cuticle thickness and cuticle permeation thickness.

### Cuticle composition analysis by Gas Chromatography-Mass Spectrometry

Fruit cuticles were isolated from wild-type, *cr-slhdziv7* and *cr-slhdziv9* mutant fruits at 20 DPA and 50 DPA. Fruit discs with a diameter of 50 mm were excised from the fruits used for cuticle extraction. Each sample consisted of pericarp tissues from five individual fruits, with three biological replicates performed for each group. The discs were incubated in a solution containing 2% cellulase and 1% pectinase at 42 °C for 3 days to remove the exocarp cells on the cuticular membranes according to the method described previously (Fich et al., 2020; Hovav et al., 2007). Subsequently, the digested tissues were rinsed off from the cuticles with distilled water.

The dried, isolated cuticular discs were sequentially soaked in chloroform and methanol to remove wax components. The dewaxed cuticles were then depolymerized using a mixture of methanol, sodium methoxide and ethyl acetate, with the addition of 20 μg of *n*-tetracosane added as the internal standard. The samples were incubated at 60 °C overnight and then cooled to room temperature. Cuticular components were extracted with dichloromethane, and the organic phase was dried under a stream of nitrogen gas. The resulting cuticular extracts were resuspended in pyridine (EMD Millipore) and derivatized with BSTFA (*N,O*-bis(trimethylsilyl)trifluoroacetamide) at 70 °C for 30 min (Fich et al., 2020). Thereafter, the samples were analyzed using a gas chromatography-mass spectrometer equipped with thermal desorption/headspace solid-phase microextraction (Agilent 7890B-5977B, USA) under previously established parameters (Martin et al., 2017). The oven temperature was held at 60 °C for 1 min, then increased at a rate of 5 °C per minute up to 300 °C and held at this temperature for 10 min to achieve the separation of cutin monomers. Compound abundance was determined by integrating the chromatographic peaks using ChemStation software and normalizing the peak areas against the internal standard. Compounds were identified by comparing mass spectra with the NIST Mass Spectral library (Version 2.2).

### Gene expression analyses

Total RNA was extracted from the fruit trichomes and pericarp tissues of wild-type, *cr-slhdziv7* and *cr-slhdziv9* mutants at 20 DPA, where the plant tissues in each sample were pooled from three individual fruits. The fruits were harvested and stored immediately at −80 °C for 24 hours. For fruit trichome isolation, the frozen fruits were placed on precooled watch glasses positioned above liquid nitrogen (with no contact with the liquid nitrogen) to maintain cryogenic conditions throughout the procedure. The fruit trichomes were carefully brushed from the surface of the fruit using a standard paintbrush that had been precooled by immersion in liquid nitrogen. The detached trichomes were collected in the watch glasses and transferred to sterile microcentrifuge tubes and stored at −80°C until RNA extraction. RNA extraction was performed using the Plant Total RNA Extraction Kit (Tiangen, China) according to the manufacturer’s instructions. First-strand cDNA was synthesized from the extracted RNA using the HiScript IV All-in-One Ultra RT SuperMix for qPCR Kit (Vazyme, China). For quantitative real-time polymerase chain reaction (RT-qPCR) analysis, the obtained cDNA templates were diluted 50-fold. Reactions were prepared using the SuperReal PreMix (SYBR Green) Kit (Tiangen, China) and performed on a LightCycler® 96 System (Roche, Switzerland) with a three-step PCR protocol. The relative expression levels were calculated using the 2^□ΔΔCt^ method, with *SlRPL2* was selected as the internal reference. Each assay was conducted with three technical replicates. The primers are listed in Table S1.

The RNA sequencing data from the previous study (Li et al., 2025) were deposited in the National Center for Biotechnology Information (NCBI) Sequence Read Archive under the accession number PRJNA1377223. RNA was extracted from three-week-old leaves of the *cr-slhdziv9* mutants and WT plants. Annoroad Gene Technology (Beijing) performed the library preparation and sequencing on an Illumina NovaSeq X Plus platform to generate paired-end reads. The adaptor sequences, reads containing ploy-N, and low-quality sequence reads were removed from the datasets. Clean reads were subsequently mapped to the *Solanum lycopersicum* ITAG4.0 reference genome using HISAT2 v2.2.1 software (Kim et al., 2019). The expression level of each gene was normalized according to the transcripts per million reads (TPM) method. The DESeq2 R package was used to identify differentially expressed genes (DEGs), with thresholds set at |log2FoldChange| > 1 and an adjusted *P*-value of < 0.05.

### Postharvest water loss assays

At least 20 uniformly sized fruits at the full-ripe stage were sampled from each line (WT, *cr-slhdziv7*, and *cr-slhdziv9*). The pedicel abscission zone was sealed with paraffin. Each line of ten fruits was carefully wiped off the trichomes from the surface with absorbent paper to simulate damage to fruit trichomes during harvesting and transportation. After that, all the fruits from different groups were placed at room temperature with 40-50% relative humidity, and their fresh weights were measured every two days for 20 consecutive days. The fruit water loss percentage of each group was calculated according to the following formula: Fruit water loss (%)= (W_0_-W_n_)/W_0_ ×100, ‘W_0_’ is initial fruit weight, ‘W_n_’ represents the weight after n days of storage.

### Yeast One**□**Hybrid Assays

To investigate the potential interaction between the transcription factors SlHDZIV7 and SlHDZIV9 and the *SlCYP86A68* promoter, a yeast one-hybrid (Y1H) assay was performed. The coding sequences of SlHDZIV7 and SlHDZIV9 were cloned into the pB42AD prey vector (Clontech), generating pB42AD-SlHDZIV9 and pB42AD-SlHDZIV7. The 1-2 kb fragments of the *SlCYP86A68* promoter were amplified and inserted into the pLacZ-2u reporter vector (Clontech) to generate pLacZ-2u-*proSlCYP86A68*.The primers used are listed in Table S1.

Pairs of recombinant vectors were co-transformed into the yeast strain EGY48 (Clontech) using the lithium acetate method. Positive transformants were selected on synthetic dropout (SD) medium SD/-Ura/-Trp agar plates at 30°C for 3–4 days. Interactions were validated by β-galactosidase activity assays using X-gal, with blue colonies indicating TF-promoter interactions. The experiments were performed three times with similar results.

### Yeast Two**□**Hybrid Assays

To investigate potential protein-protein interactions between SlHDZIV9/SlHDZIV7 and SlMIXTA-like, a yeast two-hybrid (Y2H) assay was conducted. Yeast two-hybrid assays were performed following the Invitrogen ProQuest Two-Hybrid System instructions with modifications as previously described (Shen et al. 2016). The full-length coding sequences of SlHDZIV7, SlHDZIV9 and SlMIXTA-like were first cloned and inserted into pENTR-TOPO vector. SlHDZIV7 and SlHDZIV9 were then recombined into the pDEST32 bait vector (Invitrogen) using Gateway® LR Clonase II reaction, yielding pDEST32-SlHDZIV7 and pDEST32-SlHDZIV9. The SlMIXTA-like was recombined into the pDEST22 prey vector (Invitrogen) to generate pDEST22-SlMIXTA-like via LR reaction. The bait (pDEST32 derivatives) and prey (pDEST22-SlMIXTA-like) constructs were co-transformed into yeast strain AH109 (Clontech) using the lithium acetate method. Transformants were selected on synthetic defined (SD) medium lacking leucine and tryptophan (SD/-Leu/-Trp) agar plates at 30°C for 3–4 days. Positive colonies were further screened on high-stringency SD medium (SD/-Ade/-His/-Leu/-Trp). The experiments were repeated twice. The primers are listed in Table S1.

### Bimolecular Fluorescence Complementation (BiFC)

The Gateway-compatible BiFC vectors pEarleyagte201-YN and pEarleygate202-YC were used for BiFC analysis as previously described (Shen et al. 2016). The full-length coding sequences of SlHDZIV9 and SlMIXTA-like were first cloned and inserted into pENTR-TOPO vector. Gateway cloning was performed using LR Clonase II enzyme mix (Thermo Scientific) to shuttle the gene of interest from gateway entry clones into the destination vector (pEarleyagte201-YN and pEarleygate202-YC), following the manufacturer’s standard protocol. SlHDZIV9 was fused in-frame with EarleyGate 202-YC (C terminus of YFP) whereas SlMIXTA-like was fused in-frame with pEarleyGate 201-YN (N terminus of YFP). All vectors were then transformed into *A. tumefaciens* strain GV3101. Binary vector combinations were co-delivered into *N. benthamiana* leaves via *Agrobacterium*-facilitated infiltration. Fluorescence microscopy performed on the FV3000 confocal laser scanning microscope platform (Olympus). The YFP signals were observed with argon laser excitation at 488 nm and a 505–550 nm emission filter set. The primers are listed in Table S1.

### Dual-luciferase assays

The dual-luciferase assays were performed as described previously (Li et al., 2025). Full length of SlHDZIV9 and SlMIXTA-like coding sequences were cloned and inserted into pHB vector as the effector constructs, and the promoter fragments of target genes were cloned into pGreenII 0800-LUC vector as the reporter constructs. All vectors were transformed into *A. tumefaciens* strain GV3101, and the pHB empty vector was used as a negative control. The reporters and effectors were mixed in a 1:2 ratio and further transiently co-expressed into 4-week-old *Nicotianaben thamiana* leaves. The LUC and REN fluorescence values were measured using a commercial Dual Luciferase Reporter Gene Assay Kit (YEASEN, China) on the fluorescence detector (Promega, GloMax 20/20). At least five independent biological repeats were performed for each combination. The primers are listed in Table S1.

### Qualification and Statistical analysis

GraphPad Prism 8 software (Version 2020; USA) were used for statistical analyses. All data are presented as mean ± standard error (SE). Sample size (n) indicates biological replicates. Statistical significance was assessed by Two-tailed Student’s t-test for pairwise comparisons. Significant differences are indicated by asterisk (\**p* < 0.05, \*\**p* < 0.01). One-way analysis of variance (ANOVA) followed by Tukey’s post hoc test was used for statistical analysis among multiple groups. A *p*-value < 0.05 was considered significant.

## Results

### The *cr-slhdziv7* and *cr-slhdziv9* Mutants Exhibit Significantly Reduced Fruit Trichome Density

We used the mutant materials *cr-slhdziv7* and *cr-slhdziv9*, which were generated previously (Li et al., 2025a, b), for subsequent analysis to evaluate the effects of fruit trichome density and fruit cuticle on the postharvest water loss process. Knocking out *SlHDZIV7* and *SlHDZIV9* via gene editing significantly reduced fruit trichome density in both mutant lines (Figure 1A). The expression levels of *SlHDZIV7* and *SlHDZIV9* at 20 DPA were significantly higher expressed in fruit trichome (Figure 1B). When the trichomes on the surface of the fruit were gently rubbed off to simulate damage during harvesting and handling, followed by toluidine blue staining, the number of stained spots was significantly lower in the two mutant fruits than in the wild type (Figure 1C-D). Specifically, the number of stained spots in *cr-slhdziv7* was reduced by 60.2%, and in *cr-slhdziv9*, it was reduced by 47.8% compared to the wild type fruits (Figure 1E). Conversely, fruits that were handled with care and retained trichomes showed no significant difference in the stained spots between the mutants and wild-type fruits (Figure 1C-D). Those staining spots were primarily the residue bases of broken or detached fruit trichomes, as residue bases were more easily stained by Toluidine Blue (Fich et al., 2020). Therefore, the number of staining spots can be assumed to represent the number of fruits trichomes. Furthermore, scanning electron microscopy (SEM) was used to quantitatively analyze the fruit trichomes on the fruit surface. The SEM statistical results indicated that the density of fruit trichome in the mutant fruits of *cr-slhdziv7* and *cr-slhdziv9* were significantly lower than that of the wild-type fruits. Specifically, *SlHDZIV7* primarily affected trichome formation of types VI on fruit, while *SlHDZIV9* mainly impacted trichome development of type I and III on fruit (Figure1A). The results reveal that although *SlHDZIV7* and *SlHDZIV9* exhibit slight differences in regulating the formation of different types of fruit trichomes, both are positive regulators of fruit trichome formation in tomatoes. Either gene knockout results in altered fruit trichome density.

**Figure 1.**
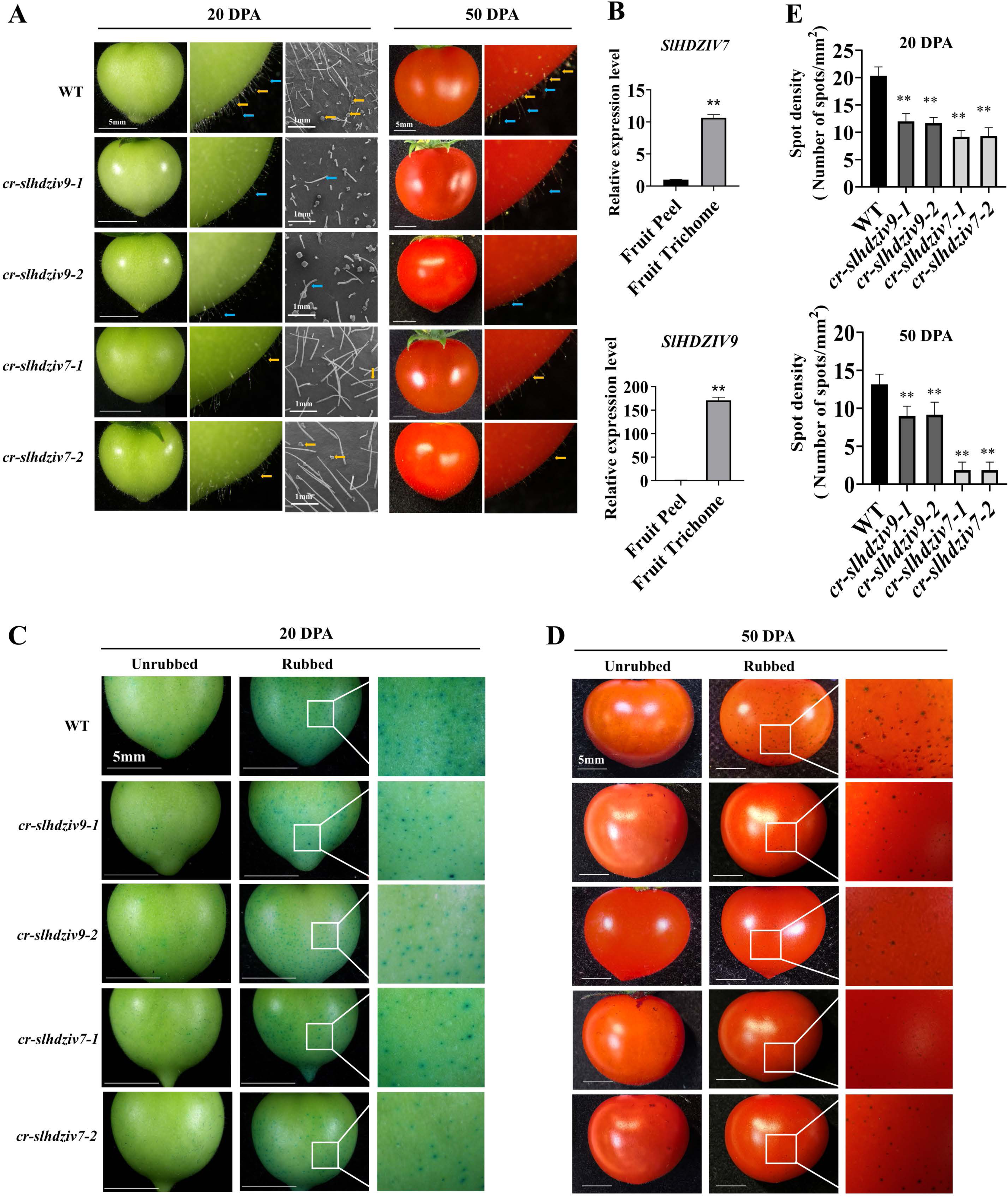
Knockout of *SlHDZIV7* and *SlHDZIV9* affects fruit trichome density and exocarp permeability. (A) Stereomicroscopy and scanning electron microscopy (SEM) reveal reduced trichome density on *cr-slhdziv7* and *cr-slhdziv9* mutant fruits compared to wild-type (WT) at 20 and 50 days post-anthesis (DPA). (B) Expression levels of *SlHDZIV7* and *SlHDZIV9* in fruit peel and fruit trichome at 20 DPA. Both *SlHDZIV7* and *SlHDZIV9* were significantly higher expressed in fruit trichome. The data represent means + SD of 3 biological replicates. Asterisks indicate statistically significant differences (Student’s *t*-test: **, *P* < 0.01). (C-E) Toluidine Blue staining assays. Fruits with intact trichomes limit dye infiltration, resulting in fewer blue spots. In contrast, rubbing the fruit trichomes significantly increases staining, indicating that microchannels at broken trichome bases facilitate permeation. In fruit trichome rubbed set, both mutants exhibited significantly reduced staining at both the 20 DPA (C, E) and 50 DPA (D, E) stages compared to the WT, demonstrating lower exocarp permeability. The blue spots represent dye entry through microchannels formed after trichome detachment. The average values and error bars were means ± SD of 5 individual fruits for each line. Asterisks indicate statistically significant differences (Student’s *t*-test: **, *P* < 0.01).

### The *cr-slhdziv7* and *cr-slhdziv9* mutations exhibit thinner cuticle thickness

Trichomes and cuticles are both key protective epidermal specializations. The genetic interplay connection between trichome and cuticle formation has been previously suggested in a variety of species (Berhin et al., 2022; Chalvin et al., 2020; Shi et al., 2017; Yan et al., 2018). Several tomato genes have also been reported to influence the formation of both trichomes and cuticles, such as *SlCD2* (Nadakuduti et al., 2012), Woolly (Xiong et al., 2020), SlMX1 (Ewas et al., 2016) and SlMIXTA-like (Galdon-Armero et al., 2020). To determine the effect of *SlHDZIV7* and *SlHDZIV9* on cuticle formation, we observed the cuticle thickness at multiple developmental stages of the mutant fruits by histological staining (Figure 2A). The thickness of the cuticle was defined either in the thinner zone at the midpoint of the upper layer of the epidermal cells (Z1) or the thicker zone at the junction of two pavement cells (Z2). A quantitative analysis of the change in cuticle thickness revealed a variation in cuticle thickness among the two types of mutants and WT fruits. Specifically, at the 20 DPA stage, the cuticle of the WT and mutant fruits were found to be relatively thin, with an average thickness of approximately 3.6-4.9 μm for Z1 point and 9.8-14.2 μm for Z2 point (Figure 2B-C). The *cr-slhdziv7* mutant exhibits significantly reduced Z1 thickness compared to the WT, whereas the *cr-slhdziv9* mutant shows no significant difference in Z1 thickness relative to the WT (Figure 2B). Unexpectedly, at 20 DPA, the Z2 cuticle thickness of both the *cr-slhdziv7* and *cr-slhdziv9* mutants was significantly higher than that of the WT (Figure 2C). In the period up to 30-40 DPA, the cuticle thickness in the Z2 of the *SlHDZIV7* and *SlHDZIV9* mutant fruits were even higher than in the wild-type (Figure 2C). The thickness of the cuticle was more developed in all fruit at the 50 DPA stage. Importantly, this stage coincides with the harvest period, and cuticle thickness was one of the factors that influenced postharvest water loss. In comparison to the wild type (6.3 ± 0.7 μm in the Z1 and 22.5 ± 2.2 μm in the Z2 point), the cuticles of the mutant types *cr-slhdziv7* and *cr-slhdziv9* exhibited thicknesses of 5.3 ± 0.4 μm and 4.9 ± 0.6 μm in the Z1 and 14.9 ± 1.4 μm and 14.8 ± 1.9 μm in the Z2 point, respectively (Figure 2B-C). The results indicate that the rate of cuticle biosynthesis during fruit development is significantly reduced by knocking out the *SlHDZIV7* or *SlHDZIV9* genes, ultimately resulting in thinner cuticles in ripening fruit. This suggests that these genes are involved in cuticle formation in addition to trichome formation.

**Figure 2.**
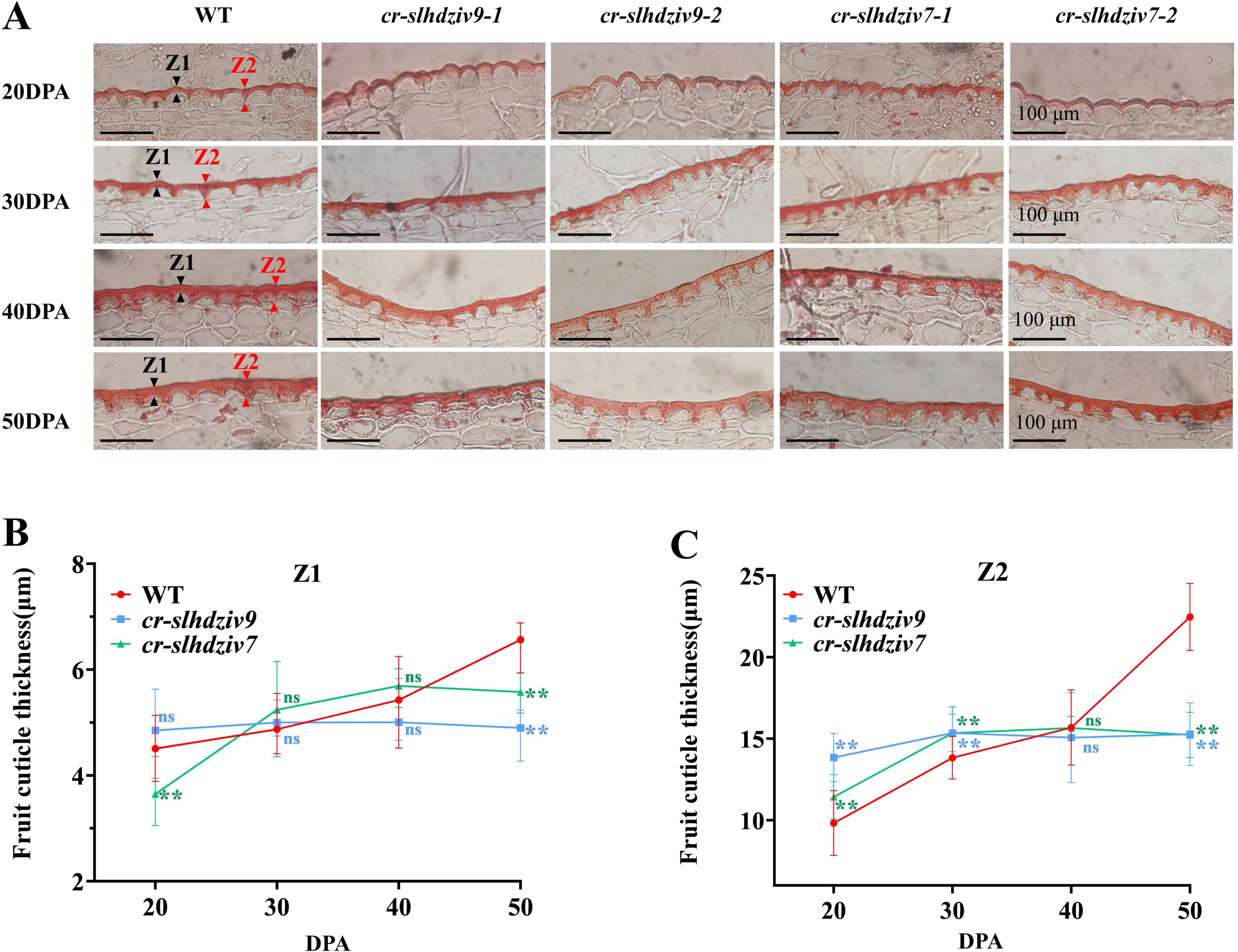
Cuticle deposition is reduced in *SlHDZIV7* and *SlHDZIV9* mutant fruits. (A) Morphology of cryosection of outer pericarps from WT, *cr-slhdziv7*, and *cr-slhdziv9* fruits at 20, 30, 40, and 50 DPA, stained with Oil Red O and observed by light microscopy. Black and red arrows indicate the cuticle thickness at the midpoint of the upper epidermal layer (Z1) and at the two pavement cells junction (Z2), respectively. Scale bars: 100 μm. (B) Cuticle thickness at Z1 point in the WT, *cr-slhdziv7*, and *cr-slhdziv9* lines at the 20, 30, 40, and 50 DPA stages. Mutants show significantly thinner cuticles than WT at 50 DPA. Data are means (±SD) of at least 20 cryosection slices from 4 biological replicates. Asterisks indicate statistically significant differences (Student’s *t*-test: **, *P* < 0.01, ns, not significant). (C) Cuticle thickness at Z2 point in the WT, *cr-slhdziv7*, and *cr-slhdziv9* lines at the 20, 30, 40, and 50 DPA stages. Mutants show significantly thinner cuticles than WT at 50 DPA. Data are means (±SD) of at least 20 cryosection slices from 4 biological replicates. Asterisks indicate statistically significant differences (Student’s *t*-test: *, *P* < 0.05; **, *P* < 0.01, ns, not significant).

### The *cr-slhdziv7* and *cr-slhdziv9* mutations altered the cuticle composition and properties in tomatoes

To examine the roles of SlHDZIV7 and SlHDZIV9 in fruit cutin biosynthesis, and to determine whether the biochemical composition of the cuticle was altered in the mutants, the fruit peels of the WT, *cr-slhdziv7* and *cr-slhdziv9* at 20 and 50 DPA were analyzed by GC-MS following standard depolymerization and derivatization procedures (Fich et al., 2020; Jin et al., 2025). The GC-MS analysis revealed that at 20 DPA, there is no significant difference between the mutant lines *cr-slhdziv7* and *cr-slhdziv9* and the wild type in terms of the total amount of cutin monomers and the content of ω-OH fatty acids (C16), which constitute the major fraction of cutin (Figure 3). The findings were in line with the results obtained from measuring cuticle thickness using section staining at 20 DPA (Figure 2B). In contrast to the immature fruit at the early fruit developmental stage at 20 DPA, the mature fruit at 50 PDA of the *cr-slhdziv7* and *cr-slhdziv9* fruits contained significantly lower total amount of cutin monomers and lower amounts of several predominant cutin monomers, including ω-OH fatty acids (16C), α, ω-dicarboxylic acids (16C), when normalized to pericarp area (Figure 3B, Figure S4). Despite the fact that studies have reported that wax constitutes only a small proportion of the cuticle in tomato fruit, its composition and content also influence postharvest water loss rates. The analysis of the expression profiles of genes associated with wax synthesis in two mutants revealed no consistent patterns, therefore the changes in wax composition within the mutants were not determined.

**Figure 3.**
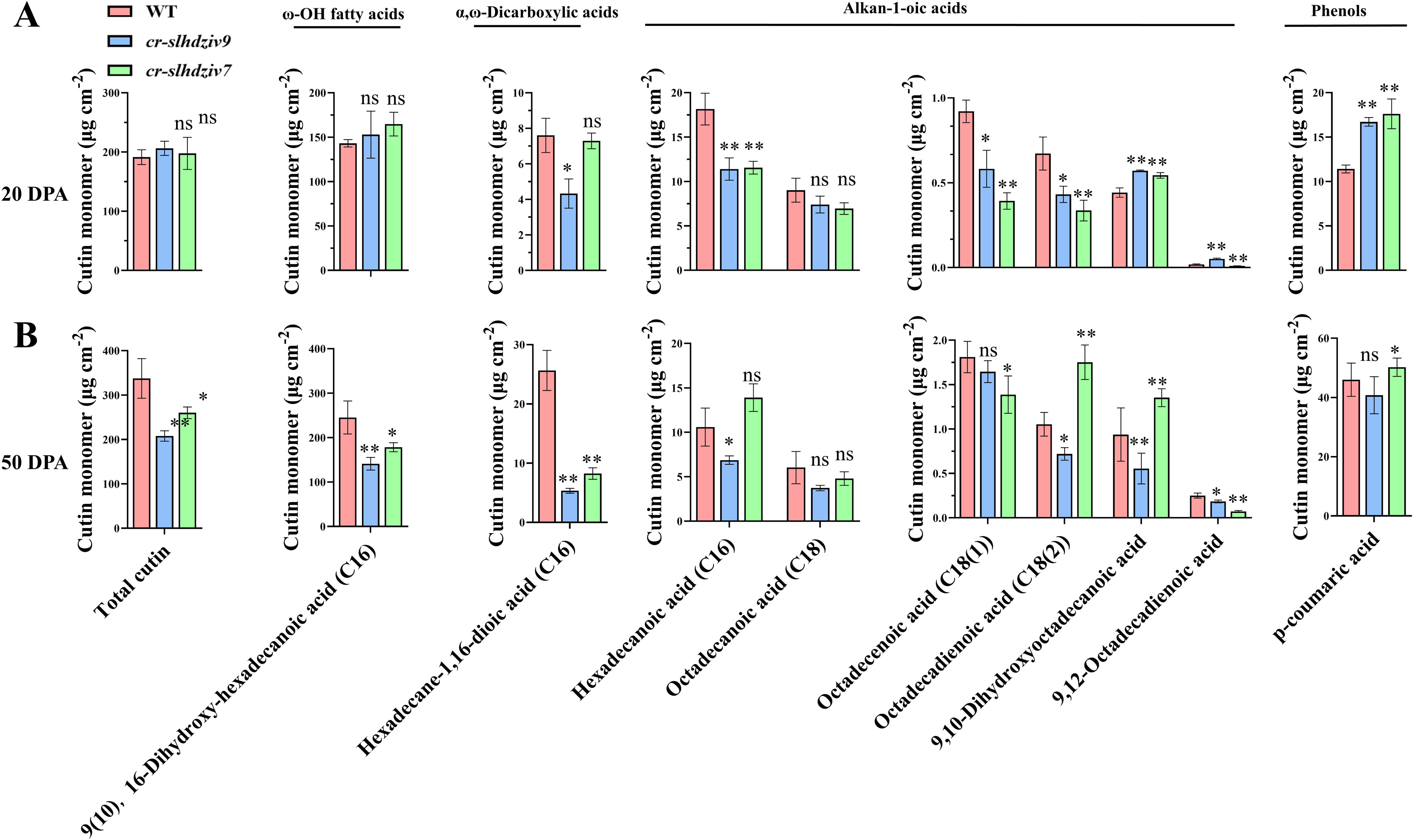
GC-MS analysis of the cutin composition in cuticles isolated from WT, *cr-slhdziv7*, and *cr-slhdziv9* lines at 20 and 50 DPA. (A) The detailed composition and contents of the WT, *cr-slhdziv7*, and *cr-slhdziv9* lines at 20 DPA stage. There is no significant difference in the total cutin monomers and predominant ω-hydroxy fatty acids between the mutants and WT at the 20 DPA stage. The data represent means (±SD) of 5 biological replicates. Asterisks indicate statistically significant differences (Student’s *t*-test: *, *P* < 0.05; **, *P* < 0.01, ns, not significant). (B) The detailed composition and contents of the WT, *cr-slhdziv7*, and *cr-slhdziv9* lines at 50 DPA stage. At 50 DPA, *cr-slhdziv7* and *cr-slhdziv9* mutants contain significantly reduced total cutin monomers and predominant ω-hydroxy fatty acids compared to WT. The data represent means (±SD) of 5 biological replicates. Asterisks indicate statistically significant differences (Student’s *t*-test: *, *P* < 0.05; **, *P* < 0.01, ns, not significant).

The GC-MS component analysis results were consistent with the histological staining results, which showed that knocking out either the *SlHDZIV7* or *SlHDZIV9* genes leads to inhibited cutin biosynthesis and results in a thinner fruit cuticle in mature fruit. These results suggest that SlHDZIV7 and SlHDZIV9 act as positive regulators of cutin biosynthesis in tomato fruit.

### SlHDZIV7 and SlHDZIV9 regulate genes involved in cutin biosynthesis

To better understand the molecular basis of the changes in the composition of fruit cutin resulting from *cr-slhdziv7* and *cr-slhdziv9*, we measured changes in the expression of cutin biosynthesis-associated genes. In previous studies on the role of the *SlHDZIV9* gene in regulating trichome formation, we completed the transcriptomic sequencing analysis of *cr-slhdziv9* and WT in leaves. Although we did not perform RNA-seq on fruit peel, the epidermal layer of leaves is also covered by cuticle, and Toluidine Blue staining analysis of leaf permeability also revealed that the *cr-slhdziv9* exhibited thinner leaf cuticles and higher leaf permeability (Figure S1). Therefore, we hypothesized that the transcriptomic sequencing data from the *cr-slhdziv9* leaf sample remains valuable. To characterize expression changes in the *cr-slhdziv9* and WT samples, we conducted the DEGs analysis, which were identified using the following criteria: |log2FoldChange| ≥ 1 and FDR ≤ 0.05. There were 419 DEGs identified in the *cr-slhdziv9* vs WT (Figure S2). To interpret the molecular mechanisms of the DEGs, the Kyoto Encyclopedia of Genes and Genomes (KEGG) pathway enrichment analysis was performed using those DEGs. The results revealed that the DEGs were enriched in pathways such as plant secondary metabolism, cutin and fatty acid biosynthesis (Figure S3). Those indicated that the SlHDZIV9 transcription factor likely regulates genes related to the cutin biosynthetic pathway. To further ensure the accuracy of the data, we verified the expression level of genes related to the cutin biosynthetic pathway in the peel of *cr-slhdziv7*, *cr-slhdziv9* mutants and WT fruits at 20 DPA, using qRT-PCR. The results showed that the expression of several genes involved in cutin synthesis was significantly altered in *cr-slhdziv7* and *cr-slhdziv9*, such as *SlCYP86A68* (Figure 4B).

**Figure 4.**
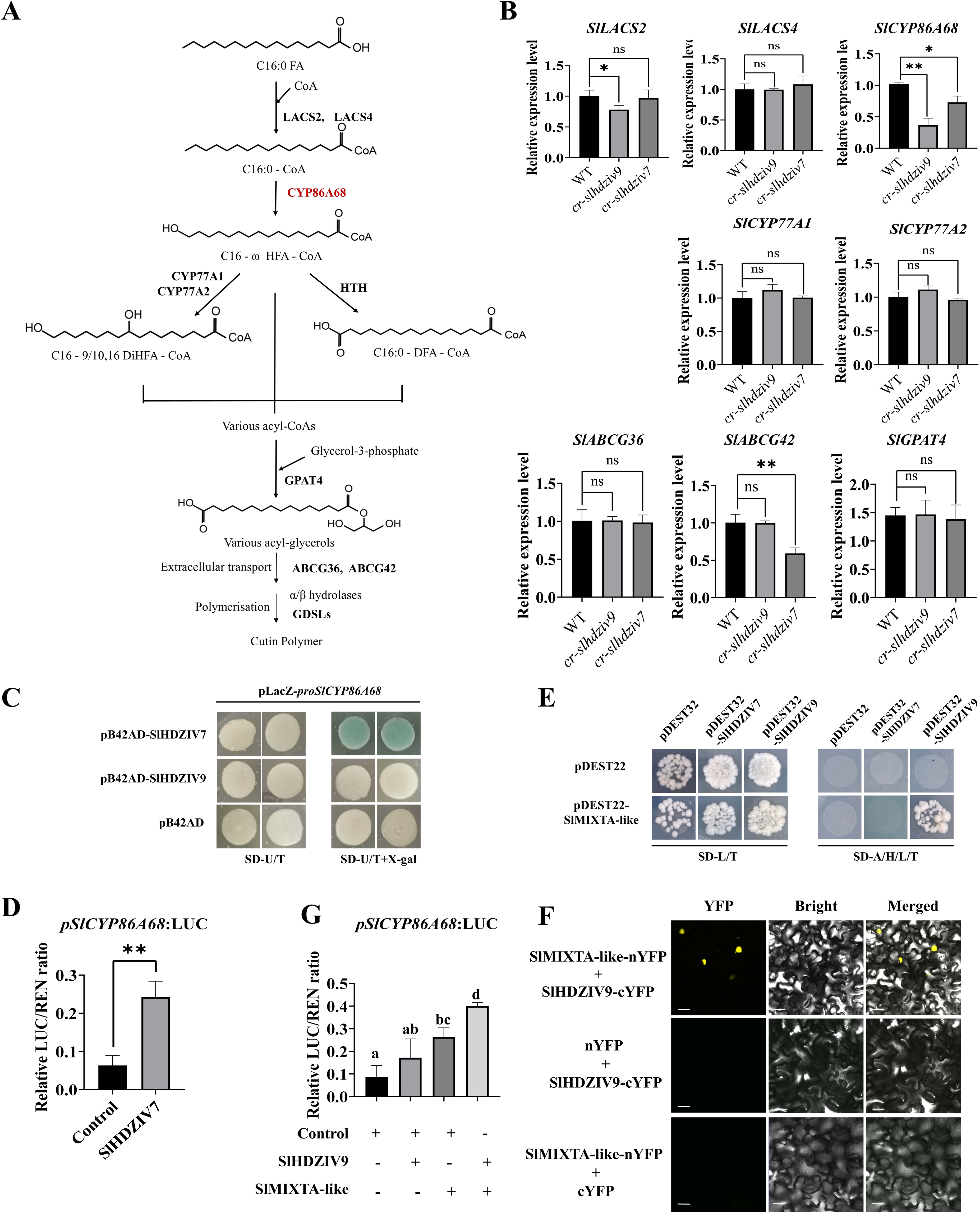
SlHDZIV7 and SlHDZIV9 transcription factors regulate genes involved in cutin biosynthesis. (A) Simplified schematic of cutin biosynthetic pathway. (B) Expression levels of cutin biosynthesis pathway genes in pericarps of 20 DPA fruits. The relative expression levels of genes were compared between the WT and *cr-slhdziv7*, and *cr-slhdziv9* lines. The data represent means + SD of 3 biological replicates. Asterisks indicate statistically significant differences (Student’s *t*-test: *, *P* < 0.05; **, *P* < 0.01, ns, not significant). (C-D) Yeast one-hybrid (Y1H) and dual-luciferase (Dual-LUC) assays demonstrate SlHDZIV7 binding and transcriptional activation of *SlCYP86A68* gene. The data represent means + SD of 5 biological replicates. Asterisks indicate statistically significant differences (Student’s *t*-test: **, *P* < 0.01). (E-F) Yeast two-hybrid (Y2H) and bimolecular fluorescence complementation (BiFC) experiments confirm the interaction between SlHDZIV9 and SlMIXTA-like, whereas no interaction was detected between SlHDZIV7 and SlMIXTA-like. (G) Dual-LUC assays show the SlHDZIV9/SlMIXTA-like complex enhances SlMIXTA-like-mediated activation of *SlCYP86A68* gene. Data represent means ± SD of 5 biological replicates. One-way analysis of variance (ANOVA) followed by Tukey’s post hoc test was used for statistical analysis among multiple groups. A *p*-value < 0.05 was considered significant.

To investigate whether SlHDZIV7 and SlHDZIV9 proteins directly regulate these cutin synthesis genes, yeast one-hybrid (Y1H) and dual-luciferase reporter assays were performed. The results indicated that the SlHDZIV7 transcription factor can bind to and activate the promoter regions of the *SlCYP86A68* genes, key gene in cutin biosynthesis (Figure 4C, D). However, SlHDZIV9 failed to bind to the promoter regions during the Y1H assay (Figure 4C). Previous studies have reported that the plant MIXTA transcription factors play key role in regulating cutin synthesis genes, and this regulatory function is conserved across multiple plant species such as Tomato, Arabidopsis, Sweet Wormwood (Galdon-Armero et al., 2020; Oshima et al., 2013; Shi et al., 2018). Additionally, the MIXTA type transcription factors often interact with the HD-ZIP IV transcription factor (Wu et al., 2018; Yan et al., 2018; Yang et al., 2022). We hypothesize that SlHDZIV9 may regulate the expression of cutin synthetic pathway genes by interacting with SlMIXTA. Notably, Y2H and BiFC experiments demonstrated that SlHDZIV9 can interact with SlMIXTA-like, while SlHDZIV7 demonstrates no such interaction (Figure 4E, F). Furthermore, Dual-LUC experiments revealed that SlHDZIV9, by interacting with SlMIXTA-like, can enhance SlMIXTA-like ability to activate transcription of downstream genes *SlCYP86A68* (Figure 4G). We speculate that SlHDZIV9 may participate in cuticle synthesis indirectly. These findings suggest that SlHDZIV7 and SlHDZIV9 regulate cutin biosynthesis genes directly and indirectly.

### Reduced fruit trichome density overrides the thin cuticle and significantly decreases postharvest water loss

To evaluate the functional impact of altered fruit surface morphology, standardized postharvest water loss assays were performed. Fruits at the red ripe stage of *cr-slhdziv7*, *cr-slhdziv9*, and WT were harvested, surface-dried, and stored at 25 ± 2 °C with 40-50 % relative humidity. To simulate the damage caused to the fruit trichome during tomato harvesting or transportation, we conducted postharvest water loss experiments using artificially damaged trichomes (wiping the fruit surface gently with soft tissue). Cumulative water loss was calculated from fresh weight measurements taken on day 0 and at regular 2 days intervals for 20 days. Interestingly, loss of function of the *SlHDZIV7* or *SlHDZIV9* genes resulted in a thinner and lower-composition cuticle layer. However, these two mutants exhibited lower water loss rates (Figure 5A). During the experimental period, at day 6, the water loss rate of *cr-slhdziv7* and *cr-slhdziv9* mutants were significantly lower than that of WT (Figure 5B). Correspondingly, external quality traits such as reduced shriveling indicated extended shelf life for the mutants under the tested conditions (Figure 5A). However, if the fruit trichomes remain intact, the postharvest water loss rates of *cr-slhdziv7* and *cr-slhdziv9* mutants show no significant difference compared to WT fruit (Figure S6). The results of the shelf life analysis were consistent with those of the fruit permeability assay via TB staining. The TB staining assay showed that *cr-slhdziv7* and *cr-slhdziv9* mutant fruits stained less intensely than WT fruits, indicating significantly lower epidermal permeability and a more effective water barrier (Figure 1C, D). Consequently, despite producing cuticle thinning, it ultimately results in lower postharvest transpiration and an extended postharvest shelf life. Together with the observed decrease in trichome density, expected to reduce trichome-associated microchannels that facilitate transpiration, these findings support the conclusion that reduced trichome abundance improves the overall barrier to water loss in the mutants, overriding the water-permeability effects typically associated with a thinner, compositionally lighter cuticle.

**Figure 5.**
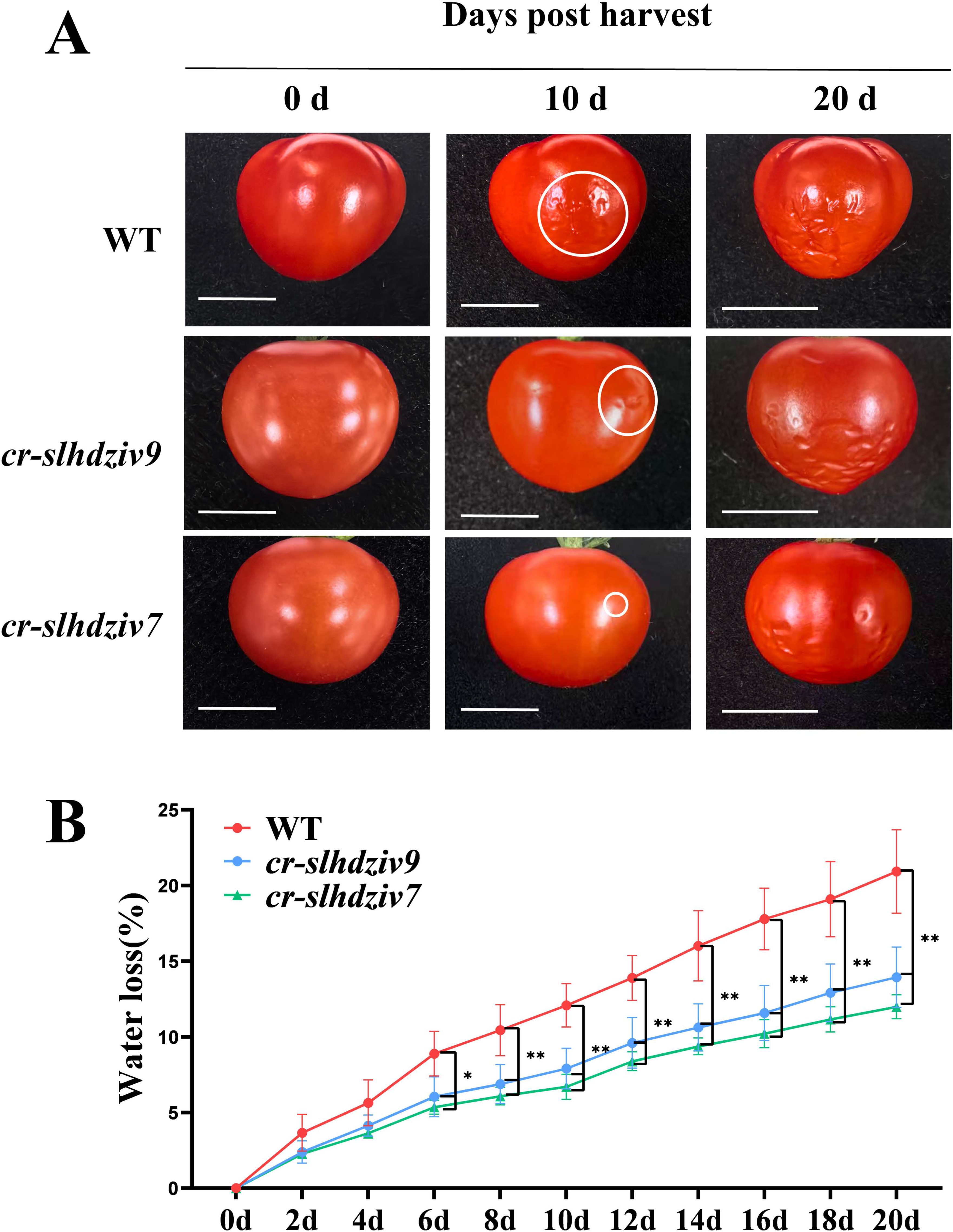
Knockout of SlHDZIV7 and SlHDZIV9 extends the postharvest shelf life of fruits. (A) The morphology of red ripe fruits from the WT, *cr-slhdziv7* and *cr-slhdziv9* lines were examined after being stored at room temperature for 0, 10 and 20 days. Both mutant fruits exhibit prolonged shelf life. The *cr-slhdziv7* fruit shows less severe pericarp shrinking (indicated by white circle) than the *cr-slhdziv9* fruit, correlating with its lower fruit trichome density. (B) Water loss rates of fruits from the WT, *cr-slhdziv7* and *cr-slhdziv9* lines in (A). The weight of per fruit was measured every 2 days over 20 days of storage. By day 6 postharvest, both mutants display significantly reduced water loss compared to WT. Data represents the means + SD of 5 biological replicates. Asterisks indicate statistically significant differences (Student’s *t*-test: *, *P* < 0.05; **, *P* < 0.01).

## Discussion

### Fruit trichome density precedes cuticle thickness as a postharvest water barrier

In this study, we used multiple approaches to analyze the roles of trichome density and cuticle thickness in postharvest water loss in fruit. We demonstrated that two tomato mutants, *cr-slhdziv7* and *cr-slhdziv9*, exhibited reduced fruit trichome density by 60.2% and 47.8%, respectively, relative to WT (Figure 1). However, the mutant plants exhibited reduced displayed lower postharvest water loss rates and extended shelf life than WT (Figure 5), despite having thinner cuticles with reduced amounts of major cutin monomers (Figure 2, Figure 3). This seemingly paradoxical result provides a new explanation for the longstanding “cuticle paradox”, wherein cuticle-deficient mutants do not always exhibit the expected increase in water transpiration.

Prior work has implicated fruit trichome-associated microchannels function as low-resistance pathways for water diffusion. Fich et al. (2020) provided pioneering evidence that trichome bases expose structural pores that facilitate transpiration in multiple tomato cultivars. However, that study compared trichome density in cultivars with varying genetic backgrounds, made it still challenging to determine the specific impact of trichome density on water loss. More recently, Fu et al. (2026) demonstrated through gene editing to knock out *SlHDZIV14*, a key regulator gene in fruit trichome formation, that reducing fruit trichome density is sufficient to lower water loss rates, providing the first genetic validation of trichome density as a determinant of postharvest transpiration. Nevertheless, since cuticle thickness was not significantly altered in *cr-slhdziv14* lines, that study was unable to compare the roles of trichome density and cuticle properties act independently or hierarchically to control water loss.

The present study directly addresses this gap by providing the integrated in which both fruit trichome density and cuticle properties are simultaneously altered within the same genetic background. Our results demonstrate that the advantages of reducing trichome-associated microchannels appear to outweigh the negative effects of cuticle thinning in epidermal permeability (Figure 1C, D). We therefore propose a hierarchy of barriers within the complex peel system of postharvest fruit water loss, where the influence of trichome density on water loss may be more significant than the changes in cuticle thickness or composition. These findings highlight the importance of considering additional epidermal surface features when evaluating postharvest water loss in tomato fruits.

### A new strategy for postharvest water retention should focus on integrating fruit trichomes and cuticles

Our results supported a revised model for postharvest water retention which, beyond the traditional focus on cuticle thickness or composition, incorporated fruit trichome density as a core consideration (Figure 6). In this model, the cuticle still serves as a foundational barrier, and its thickness and composition remain crucial. However, they are not the sole determinants of water loss. The damage or dislodgement of fruit trichomes were among the main factors affecting cuticle layer integrity. A thinner but structurally intact cuticle layer might perform better than a thicker, structurally compromised cuticle layer. This integrated model provides a scientific framework for reinterpreting prior observations in cuticle-deficient mutants. The *shn2* mutant showed severe cutin-deficiency, corresponding to a reduction of over 80% compared with the WT and presented a smooth and glossy fruit surface phenotype (Bres et al., 2022), which may reflect a reduction in surface trichomes. Surprisingly, the postharvest water loss was not increased but greatly reduced in *shn2* fruit compared to the WT (Bres et al., 2022), suggesting that a similar trichome-associated mechanism could explain its unexpected water loss phenotype. However, trichome density in *shn2* has not been directly quantified and this interpretation remains inferential. Future studies directly measuring fruit trichome density in *shn2* and other cuticle-deficient mutants would help establish the generality of the trichome-cuticle hierarchy model. Together, these findings indicate that fruit trichome density and dynamics are important epidermal traits that should be systematically evaluated in studies of cuticle function and fruit postharvest physiology.

**Figure 6.**
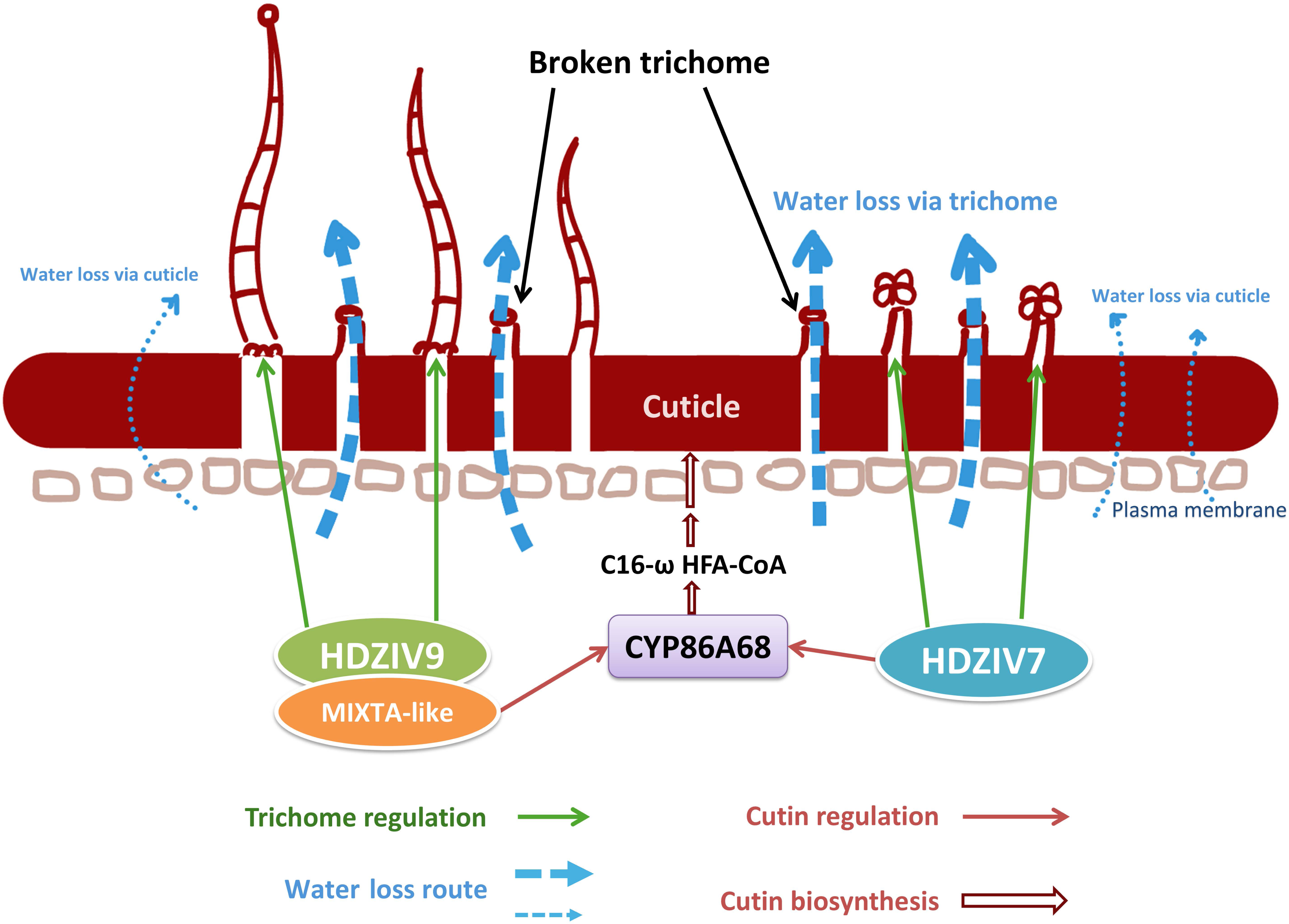
Tomato fruit trichome-associated microchannels dominate postharvest water flux. *SlHDZIV7* and *SlHDZIV9* coordinately regulate fruit trichome formation: *SlHDZIV7* primarily regulates Type VI trichomes, whereas *SlHDZIV9* controls Type I-III trichomes. Both transcription factors influence cuticle development. SlHDZIV7 directly activates *SlCYP86A68*, while SlHDZIV9 enhances the transactivation of *SlCYP86A68* by SlMIXTA-like via a protein-protein interaction. Microchannels at broken trichome bases constitute primary water loss route. Despite exhibiting thinner cuticles at 50 DPA, both mutants display significantly reduced postharvest water loss due to their lower trichome density and consequently fewer microchannels.

However, the trichome-cuticle hierarchy model proposed here was established and validated exclusively in the “Micro-Tom” genetic background. Although “Micro-Tom” provides a well-controlled genetic system ideal for investigating gene function and trichome formation, it is not yet clear whether the dominant role of trichome density over cuticle thickness in determining postharvest water loss extends to commercial tomato cultivars. Further research is needed to examine the relationship between trichome density and cuticle thickness across different genetic backgrounds. This will establish the broader applicability of the hierarchical model and evaluate its relevance for practical breeding strategies.

### Implications for Crop Improvement

In this study, we demonstrated that tomato fruit trichome-associated microchannels were the main determinants of postharvest transpiration and suggested that selectively reducing fruit trichome density represents a promising strategy for extending shelf life. Beyond tomato, broader considerations of fruit trichome density and surface architecture are relevant to postharvest quality management in diverse crops. In cucumber, the spines are a specialized form of fruit trichome. Breeding for spine-reduced or spineless varieties has resulted in smoother surfaces, which facilitates packaging, cleaning, and reducing potential pesticide residues (Yang et al., 2019). While the direct contribution of spine density to postharvest water loss in cucumber has not yet been established, it is worthy of future investigation given the structural and functional parallels with tomato trichome microchannels. In peach, studies have shown that reducing the density of peach fruit trichomes can be effective in minimizing allergies triggered by these trichomes and reducing the residual presence of pathogens (Shen et al., 2023). Despite the aforementioned advantages, it is important to acknowledge the functions of plant trichomes, including their role in defending against pests and pathogens and regulating surface microclimate (An et al., 2023; Hao et al., 2024; Jin et al., 2025; Vendemiatti et al., 2024). The breeding challenge should therefore target the fruit trichome-specific genes that regulate trichome formation, with the aim of reducing fruit trichome density while maintaining minimal impact on trichome density in other plant parts. This approach considers the benefits of reducing fruit water loss while also anticipating potential trade-offs in pest resistance or quality attributes.

## Accession numbers

Sequence data from this article are available in Solanaceae Genomics Network (https://solgenomics.net/), Phytozome 13 (https://phytozome-next.jgi.doe.gov/). The tomato genes are under the following accession numbers: *SlHDZIV7* (Solyc10g005330); *SlHDZIV9* (Solyc06g072310); *SlMIXTA-like* (Solyc02g088190); *SlLACS2* (Solyc01g109180); *SlLACS4* (Solyc01g095750); *SlCYP86A68* (Solyc01g094750); *SlCYP77A1* (Solyc01g095750); *SlCYP77A2* (Solyc01g095750); *SlABCG36* (Solyc05g018510); *SlABCG42* (Solyc01g095750); *SlGPAT4* (Solyc01g095750);

## Acknowledgements

This work has benefited significantly from the scholarly input of Dr. Glenn PHILIPPE from Laboratoire de Biologie Intégrative des Modèles Marins, Sorbonne Université and Professor Yanqun XU from School of Agriculture and Biology, Shanghai Jiao Tong University, who offered specialist suggestions on enhancing the structural framework of this article. We would like to thank Professor Pengxiang FAN from the Department of Horticulture at Zhejiang University for his help with cuticle isolation and GC-MS analysis. We also thank Dr. Yu GAO from the Shanghai Jiao Tong University Instrumental Analysis Center for the technical support and assistance in GC-MS metabolites measurement.

## Author contributions

Conceptualization: Q.S.,

Methodology: XY.L., M.L.;

Investigation: XY.L., M.L., L.H.;

Data curation and formal analysis: XT.L., S.Z., W.Z., S.X., L.Z.;

Writing—original draft: Q.S., XY.L.,

Writing—review & editing: Q.S., K.T.,

Supervision: Q.S.,

Funding acquisition: Q.S.,

## Data Availability

The raw sequence data of RNA-seq has been deposited in the Sequence Read Archive in National Center for Biotechnology Information, under the accession number PRJNA1377223.

## Supplementary Files

Table S1 Primers used in this study

Supplementary Figures

Figure S1. Knockout of *SlHDZIV7* and *SlHDZIV9* affects leaf cuticle formation and increases its permeability.

TB staining of leaves shows that knockout of *SlHDZIV7* (A) and *SlHDZIV9* (B) disrupts leaf epidermal permeability, with more areas stained with blue spots in the *cr-slhdziv7* and *cr-slhdziv9* mutants than in the WT.

Figure S2. Differentially Expressed Genes (DEGs) in *the cr-slhdziv9* mutant.

Volcano plots comparing the transcriptomes of *cr-slhdziv9* mutant with the WT. X-axis and Y-axis represent log_2_FoldChange and –log_10_(P-value), respectively. The green and red dots represent downregulated DEGs with log_2_(FC) less than –1 and upregulated DEGs with log_2_(FC) >1, respectively. The gray dots represent no significant difference in transcriptomes.

Figure S3. KEGG pathway enrichment analysis of DEGs.

The size of the point represents the number of DEGs enriched to the pathway, and the color of the point represents the Log10 Q-value of the enrichment to the pathway.

Figure S4. The selected partial GC-MS chromatograms show the fruit peel cuticle at 20 and 50 DPA. The chromatographic peaks of the main cutin monomers.

Figure S5. SlHDZIV7 is involved in regulating multiple key genes associated with the biosynthesis and transport of cutin.

(A) Yeast one-hybrid (Y1H) assays demonstrate that, in addition to *SlCYP86A68*, SlHDZIV7 also binds to the promoters of *SlGPTA4*, *SlABCG36* and *SlABCG42* genes, whereas no interaction was detected between genes promoter and SlHDZIV9.

(B) Schematic diagrams showing the predicted L1-box motifs in the promoter regions of *SlCYP86A68*, *SlGPTA4*, *SlABCG36* and *SlABCG42*.

Figure S6. Water loss rates of fruits from the WT, *cr-slhdziv7* and *cr-slhdziv9* lines with intact fruit trichomes on the surface.

(A) The morphology of 50 DPA fruits from the WT, *cr-slhdziv7* and *cr-slhdziv9* lines with intact fruit trichomes on the surface was examined after being stored at room temperature for 0, 10 and 20 days. The shelf life of fruits with intact fruit trichomes from the two mutant plants showed no significant difference compared to the WT.

(B) Water loss rates of fruits from the WT, *cr-slhdziv7* and *cr-slhdziv9* lines in (A). The weight of per fruit was measured every 2 days over 20 days of storage. Data represents the means + SD of 5 biological replicates. ns indicate statistically no significant differences (Student’s *t*-test).

